# Roles of Cro_2_ and CI_2_ binding to O_*R*_ operators in controlling lysogeny stability during prophage induction

**DOI:** 10.1101/2025.01.12.632252

**Authors:** Yiyu Pang, Xue Lei, Farid Manuchehrfar, Lin Du, Jie Liang

## Abstract

The lysogeny-lysis switch in bacteriophage lambda serves as a model for understanding cell fate decisions. The molecular network controlling this switch has been explored through extensive experimental and computational studies. Yet, the specific role of protein-DNA interactions, like the binding of CI_2_ and Cro_2_ proteins to operator sites, in regulating lysogeny stability during prophage induction remains less understood. This study employs a minimalistic model and the Accurate Chemical Master Equation (ACME) method to construct detailed probability landscapes of the network’s behavior under varying conditions, such as different dissociation and CI_2_ degradation rates which simulate UV irradiation effects. Our findings indicate that Cro_2_ binding at O_*R*_3 and CI_2_ at O_*R*_1 significantly influence lysogeny stability, with the former destabilizing and the latter stabilizing it. Conversely, interactions at O_*R*_1 by Cro_2_ and O_*R*_3 by CI show minimal impact on this stability. Through the ACME approach, we could examine the network’s global behavior under conditions unapproachable by conventional stochastic simulations. This study highlights the critical roles of specific protein-DNA interactions in maintaining lysogeny and provides insight into the broader dynamics of the lysogeny-lysis switch under various physiological stresses.

## 1 Introduction

Underneath the surface of cells is a largely uncharacterized world of omnipresent biochemical interactions which encoding gene regulatory networks giving rise to complex cellular processes such as cell fate decision [1, 2], phenotype switching [3]. Gene regulatory networks are dynamical system with multiple stable states which represent distinct cell phenotypes. Although switching between alternative states can be induced by random fluctuation, the states are very stable in many cases. Protein-DNA assemblies play central roles in regulating gene expressions, therefore, there is interest in how protein-DNA interactions influence the global behavior of gene regulatory networks and if they play important roles in cellular decision making and cell states stabilization [].

The lysogeny-lysis switch in the bacteriophage lambda serves as a paradigm for studying cell fate decision. After infecting its host Escherichia coli, the phage lambda makes a decision between two modes of living, lysis and lysogeny. In the lytic mode, phage lambda generates a large number of copies of the progeny and lyses the cell, while in the lysogenic mode, phage lambda integrates its DNA into the host genome and remains latent along with the growth and division of its host for generations. [4] The spontaneous switch rate from the lysogenic state to the lytic state is less than 10^−8^ per generation [5], which means that the lysogenic state is extremely stable. When severe DNA damage (due to UV irradiation) induces the bacterial SOS response [6], it leads to switching from lysogenic mode to lytic mode. This switching process is called prophage induction.

The molecular network that controls switching between lysogenic and lytic states has been studied extensively experimentally [7–10].The key regulatory network is composed by three operators O_*R*_1, O_*R*_2 and O_*R*_3, two promoters PRM and PR and two proteins CI and Cro. In the lysogenic mode, CI protein is expressed from the lysogenic promoter PRM and binds cooperatively as a tetramer to two operators, O_*R*_1 and O_*R*_2, to repress the lytic promoter P_*R*_ and thus block transcription of the Cro gene. In the meanwhile, this tetramer binding facilitates the transcription of its own gene from P_*RM*_. In the lytic mode, Cro is expressed from P_*R*_ and prevents CI expression, presumably by virtue of its high affinity for O_*R*_3, where it can bind to repress P_*RM*_. This CI–Cro double negative feedback loop, augmented by direct CI self-positive feedback, provides stable and heritable alternative epigenetic states. Computational modeling subsequently contributes to illuminate the central role of stochasticity and predict the critical physiological characteristics of the molecular network over time. [] The first statistical thermodynamic model of prophage induction is developed by Ackers and colleagues [11] which described the dynamics of gene regulation with cooperative binding of CI. Later in 1998, in order to explain the randomness of lysogeny-lysis decision, Arkin et al. [12] generated a fully stochastic model using the stochastic simulation algorithm (Gillespie’s algorithm) [13–15] and demonstrated that stochastic fluctuation of gene expression in a complicated system can produce probabilistic outcomes by randomizing the occurrence of crucial regulatory biochemical reactions. However, though Gillespie’s algorithm is a powerful tool to simulate the dynamics of the stochastic network and has been extremely popular and applicable, it may skip simulating rare events since it mainly simulates high-probability path at most of the computing time. And since it is difficult to determine whether the simulation is extensive enough, it is hard to say if the algorithm can provide a fully account of network stochasticity. Cao et al. [16] developed a minimalistic stochastic model using a method called discrete Chemical Master Equation(dCME) and directly computed the full steady-state probability landscape of the network from its chemical master equation. In addition, the model showed the cooperativity of CI binding to O_*R*_1 and O_*R*_2 is a critical enabling factor for phage lambda to maintain the lysogenic state.

However, a key question in understanding the lysogeny-lysis switch: whether protein-DNA interactions in the network controls the stability of the lysogeny remains unclear. Additionally, how protein-DNA interactions affect the global behaviour of the system under different physiological conditions requires further understanding. To answer these questions, we need to elucidate the exact behaviour of protein-DNA assemblies in the switching from lysogenic to lytic state. Quantitative description of switching process is mathematically and computationally challenging. In this study, we employed a method later developed by Cao et al. [17] called Accurate Chemical Master Equation (ACME, bioacme.org) to calculate exact steady-state probability landscapes of the switching network. This allowed us to investigate the network behaviour under a broad range of conditions that would not be computational feasible previously. To understand how protein-DNA interactions regulate lysogeny stability, we examined how binding of Cro to O_*R*_3/O_*R*_1 and CI to O_*R*_3/O_*R*_1 affect lysogeny stability at different dissociation rates. We further examined how such effects affect the behaviour of the system under different UV irradiation. Our results show that binding of Cro to O_*R*_3 and CI to O_*R*_1 regulate lysogeny stability: the former impairs the lysogeny stability and the latter aids in the maintenance of lysogeny during induction. Besides, we find binding of Cro to O_*R*_1 and CI to O_*R*_3 have little effects on lysogeny stability.

## 2 Results

### 2.1 Probability landscapes of lysogeny-lysis switch

To explore what controls stability of lysogeny during prophage induction, we studied a minimalistic stochastic network which contains 10 species and 50 biochemical reactions (Fig 1). Details of molecular components, binding constant and reactions rates can be found in ref [16]. We assume virus is initially at a lysogeny mode as to detect the network behavior during the transition from lysogeny to lysis.

**Figure 1:**
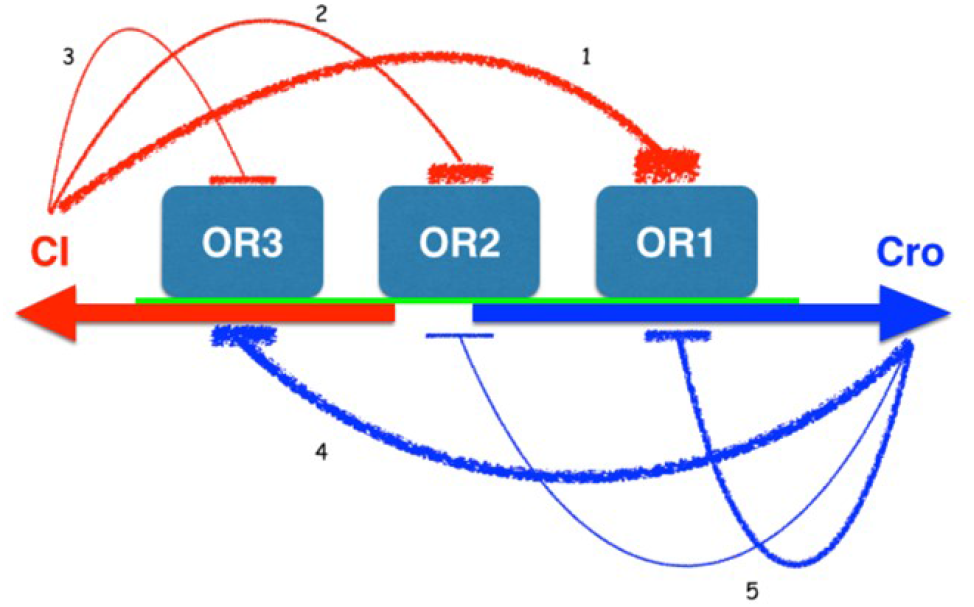
Simplified switch network

The stability of the virus states can be described by a steady-state probability landscape over the states space of the switching network, where the probability peak represents the stable states. The combination of the copy numbers of all the molecular species in the network is called the microstate of the system. Each microstate corresponds to a probability. The states space of the system is consisted of the collections of all possible microstates and the probability distribution over the states space is the probability landscape of the system. Usually, the probability landscape is evolving with time. To describe an overall behavior of the system, we focus on the steady-state probability landscape which is the result after long time evolving of probability landscape.

The switch network we studied has in total around 0.2 million microstates. Since the states space is tremendous, analytic probability landscape is challenging to be gained. The Accurate Chemical Master Equation (ACME) method we employed here cleverly solved this problem. Briefly, the method truncated the unaccessible microstates in the high-dimensional state space while the truncation error is with a predefined tolerance. Fig 2 shows the steady-state probability landscape projected to two dimensions (CI_2_ and Cro_2_), which is calculated using ACME method when degradation rate of CI_2_ is 0.0035/s. The results showed two probability peaks, where one is at high copy number of CI2 and low copy number of Cro_2_, which represents the lysogenic state, and another one with high copy number of Cro_2_ corresponding to lytic state.

Subsequently, we modeled the effects of threatening DNA damage due to UV irradiation on the network by titrating the degradation rate of CI_2_. The result(Fig 3) shows that with the degradation rate of CI_2_ increases from 0.005/s to 0.04/s, the steady-state probability landscape changes as probability peak moves from the area with high copy number of CI_2_ (lysogenic state) to the area with high copy number of Cro_2_ (lytic state). Based on the result, copy number of CI_2_ can be read as an indicator of the physiological state of the phage.

### 2.2 Cro_2_ binding to O_*R*_3 impairs lysogeny stability

To study effect of Cro_2_-O_*R*_3 binding on lysogeny stability, we varied the dissociation rate of Cro_2_-O_*R*_3 from 0.002/s to 0.009/s. Among them, 0.009/s is a wild type. We further changed the degradation rate of CI_2_ to model the process of induction. Probability landscapes of different parameter combinations was then calculated. To analyse the overall behaviour of the system, we computed the mean copy numbers of CI_2_ and Cro_2_ from the probability landscapes, where mean copy number of a protein is defined as summation of products of the protein copy number and corresponding probability at all possible microstates on a probability landscape. The results are summarised in Fig 4.

**Figure 2:**
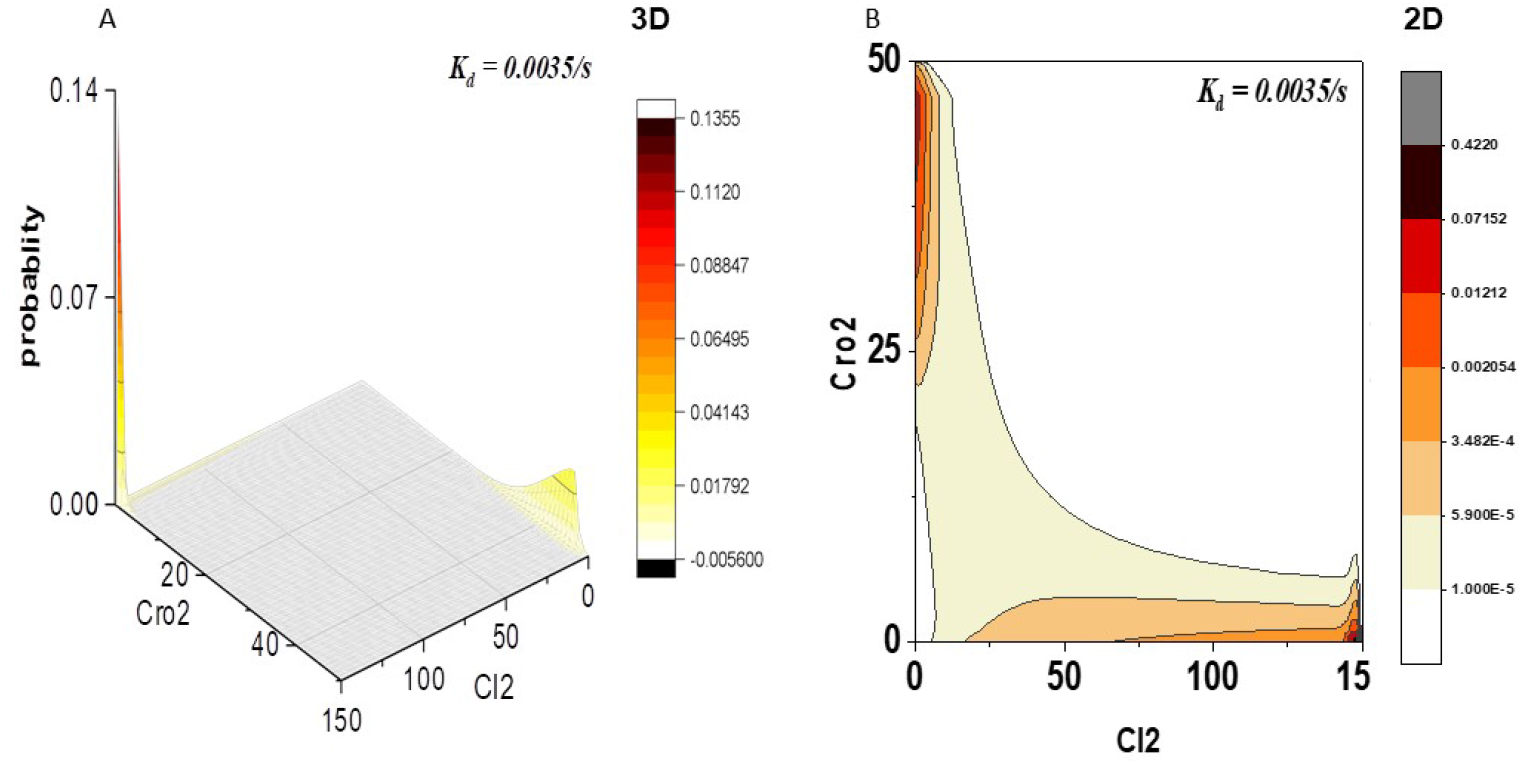
The probability landscape of switch network in phage lambda.At CI_2_ degradation rate K_*d*_ = 0.0035/s, the wild type phage lambda has two probability peak on the landscape. One is at high copy numbers of CI_2_ and Cro_2_’s copy number close to 0. Another is at high copy numbers of Cro_2_ and CI_2_’s copy number close to 0.

**Figure 3:**
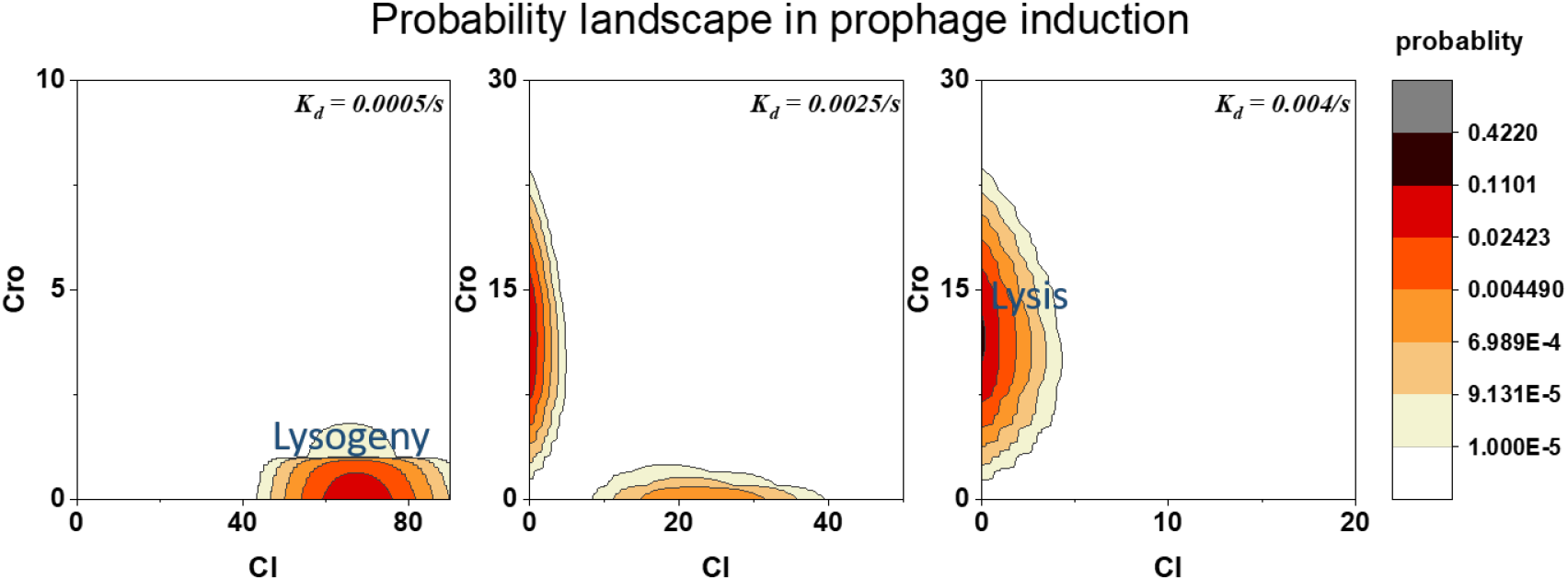
The probability landscape of switch network in prophage induction.K_*d*_ is CI_2_ degradation rate.

Our results show that binding of Cro_2_ to O_*R*_3 impairs lysogeny stability, since the switching is facilitated as the dissociation rate of Cro_2_-O_*R*_3 decreases. As in Fig 4A, for each combination of dissociation rate of Cro_2_-O_*R*_3 and CI degradation rate, we calculated the probability landscapes. we further defined the probability of lysogenic state as the total probability where CI outnumber Cro. We showed as dissociation rate of Cro_2_-O_*R*_3 decreases, the probability of lysogeny also decrease comparing at the same CI degradation rate 0.001/s. In other words, as Cro binding tighter to OR3, the switching from lysogeny to lysis is facilitated resuliting in imparing the stability of lysogeny state. Comparing the CI_2_ copy numbers’ variation(Fig 4B) of gray curve(wild type) where dissociation rate of Cro_2_-O_*R*_3 = 0.009/s and green curve where dissociation rate of Cro_2_-O_*R*_3 = 0.004/s, the CI_2_ copy number of wild type is decreasing mildly than the copy number change of green ones against changes in CI_2_ degradation rate(0.0005/s-0.002/s) due to fluctuation of UV irradiation. It shows decreasing dissociation rate of Cro_2_-O_*R*_3 causes lysogenic state unstable and facilitate the switching from lysogeny to lysis.

Our results are consistent with previous experimental observations and therefore shows the importance of Cro_2_ in the switch network. According to the work by Svenningsen et al. [18], one of their experiments which only consider the operator right region (OR) as our model did observed the derepression of PR promoter occurs at lower temperature in the cro^+^ than in cro^−^ cells in the temperature switch from lysogeny to lysis. Temperature switch models lysogeny-lysis switch by increasing temperature to inactivate CI_2_’s expression, which is similar as the increasing degradation rate of CI_2_ method in our model. Since the derepression of PR indicate the transition from lysogeny to lysis, their results suggest cro^+^ cells are induced to lytic state faster than cro^−^ cells. Therefore, our result that binding of Cro_2_ to O_*R*_3 impairs the lysogeny stability by facilitating the switch are consistent with their observation and is a good explanation to it. Schubert and Dodd’s work [19] also observed Cro_2_ binding to O_*R*_3 significantly.

**Figure 4:**
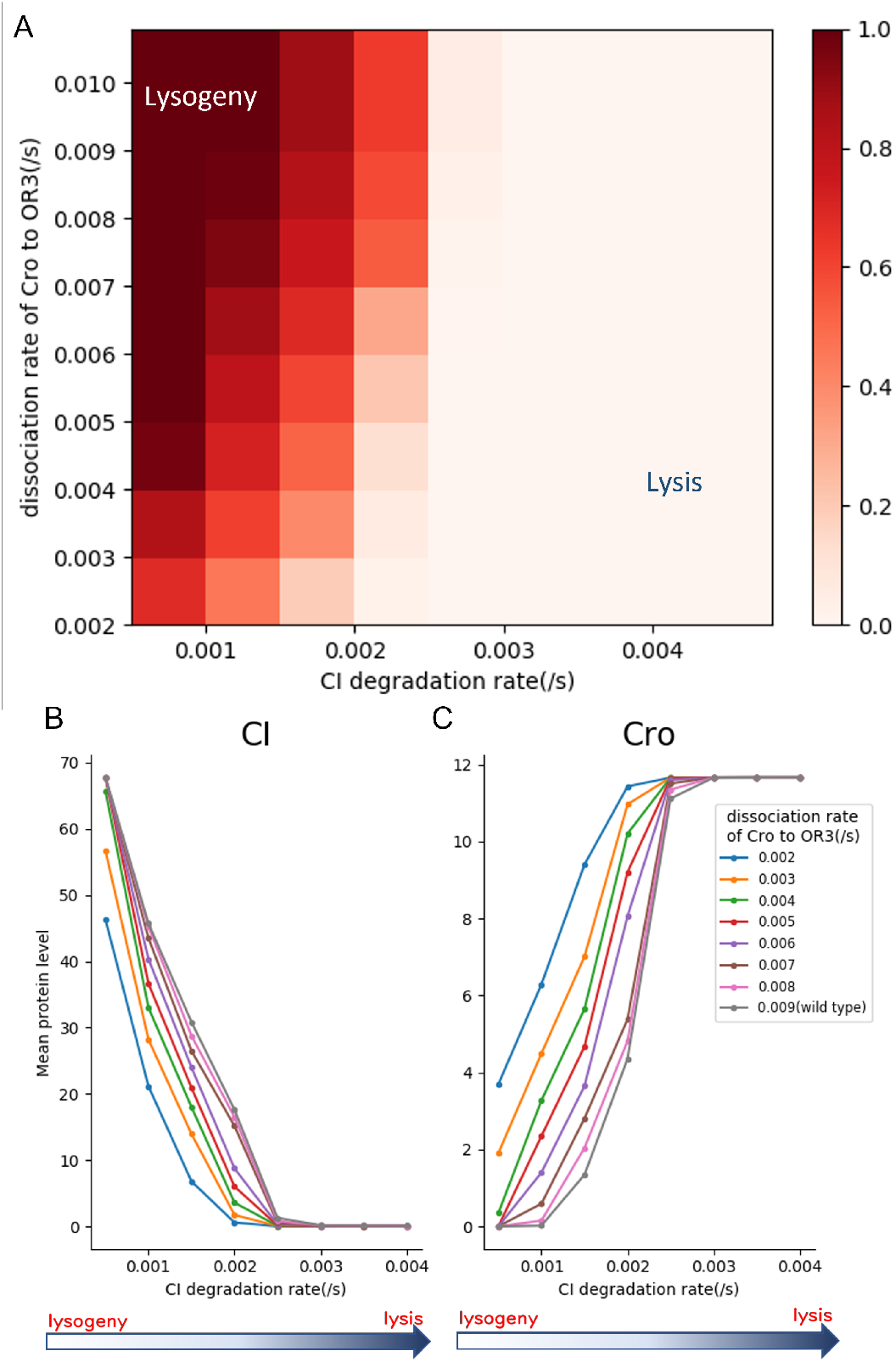
Lysogeny stability variations under different dissociation rates of Cro_2_-O_*R*_3. K_*d*_ is CI_2_ degradation rate.(A) The heatmap of lysogenic state probability under different combination of dissociation rate of Cro_2_-O_*R*_3 and CI degradation rate. (B-C) The mean copy number of CI protein(B) and Cro protein(C) in prophage induction.different color represent different dissociation rate of Cro_2_-O_*R*_3

**Figure 5:**
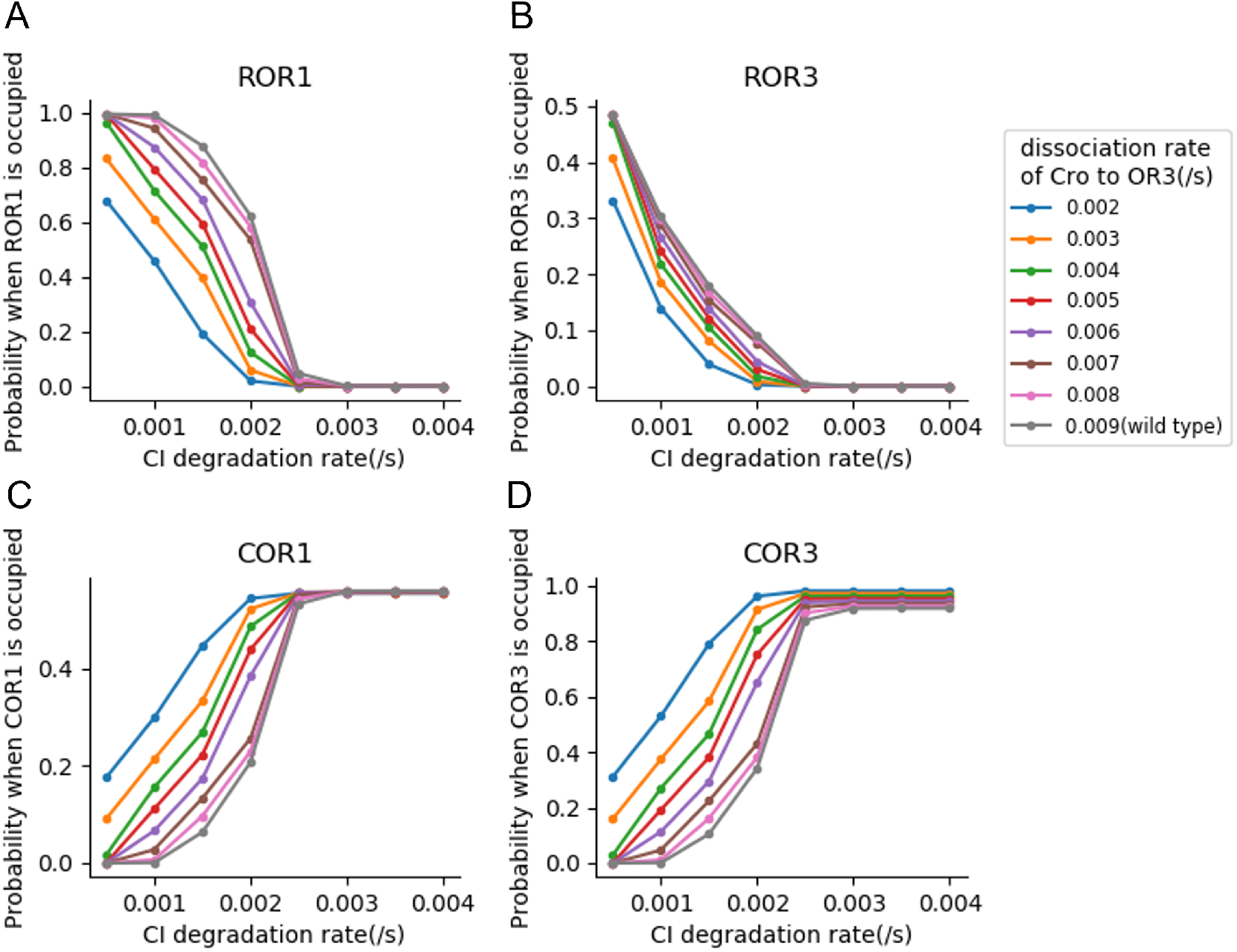
binding variation of CI(R) and Cro(C) to operator OR1 and OR3 under different dissociation rates of Cro_2_-O_*R*_3. (A) Overall probability that OR1 is occupied by CI in prophage induction. (B) Overall probability that OR3 is occupied by CI in prophage induction. (C) Overall probability that OR1 is occupied by Cro in prophage induction.(D) Overall probability that OR3 is occupied by Cro in prophage induction.

Besides, by analyzing the binding probability of CI and Cro to operate OR1 and OR3, we comfirmed the result successfully modeled the switching from lysogeny state(represented by high CI protein copy number, low Cro protein copy number) to lysis state(represented by high Cro protein copy number, low CI protein copy number). Compare Fig 5B and 5D, we show that when in lysogeny(CI degradation rate is at 0.0005/s), pRM remains active since OR3 is empty with a probability around 0.4 or occupied by CI protein with a probability of 0.3 0.5. In latter cases, synthesis of CI is promoted by binding of CI with OR3. On the other hand, Fig 5A and 5C shows in prophage the activity of pR is almost fully repressed since CI bind to OR1 with a probability nearly equals to 1. Overall, we showed that in lysogenic state, while pRM stays active, a high expression of CI lead to a nearly complete repression of pR promoter. Consistent with prior studies of lysogeny lysis switch, these results demonstrated our model correctly modeled the lysogeny state. With regarding to the lysis state (CI degradation rate is at 0.004/s), Fig 5D shows a nearly complete repression of pRM which means the synthesis of CI protein is almost blocked since Cro protein binds to OR3 with a very high probability(0.8 1). Fig 5A and 5C shows that Cro protein binds to OR1 with a probability 0.5 while CI protein doesn’t bind to OR1 at all. In conclusion, it proved the lysis state is modelled correctly, in which Cro protein can still be synthesized while the production of CI protein is blocked. In general, it can be said that our model successfully simulated the transition of lysogeny to lysis.

### 2.3 CI_2_ binding to O_*R*_1 helps maintain lysogeny stability

To study effect of CI_2_-O_*R*_1 binding on lysogeny stability, we varied the dissociation rate of CI_2_-O_*R*_3 from 0.04/s to 1.2/s. Among them, 0.04/s is a wild type. We further changed the degradation rate of CI_2_ to model the process of induction. After getting the steady state probability landscape, we calculated the mean copy number of CI_2_ and Cro_2_ at different physiological environments. We also calculated the probability of lysogeny by summing up the probability where CI outnumbers Cro.

Our results in Fig 6 show that decreasing the dissociation rate of CI_2_ from O_*R*_1 will inhibit the switch, therefore suggest that CI_2_ binding to O_*R*_1 aids to maintain lysogeny stability(Figure 5). In Fig 6A, we show as the dissociation rate of CI_2_ from O_*R*_1 decreasing, the switching happens at higher CI digradation rate where switching point is measured by the minimal needed CI degradation rate where the probability of lysogeny is nearly 0.

**Figure 6:**
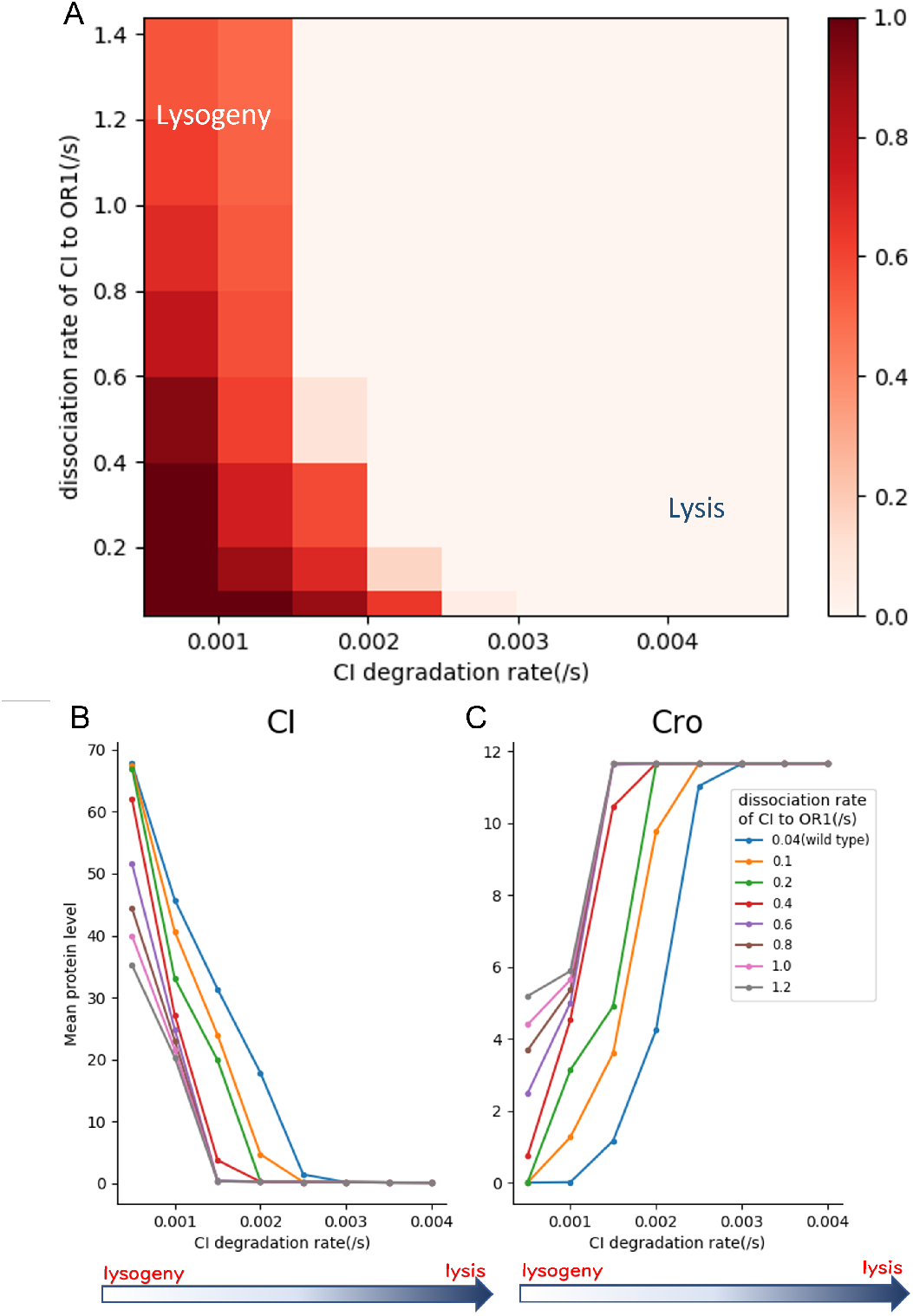
Lysogeny stability variations under different dissociation rates of CI_2_-O_*R*_1. K_*d*_ is CI_2_ degradation rate. (A) The heatmap of lysogenic state probability under different combination of dissociation rate of CI_2_-O_*R*_1 and CI degradation rate. (B-C) The mean copy number of CI protein(B) and Cro protein(C) in prophage induction.different color represent different dissociation rate of CI_2_-O_*R*_1

### 2.4 Cro_2_ binding to O_*R*_1 and CI_2_ binding to O_*R*_3 has little effect on the lysogeny-lysis switch

In order to explore the function of Cro_2_ binding to O_*R*_1 and CI_2_ binding to O_*R*_3, we varied the dissociation rates of these two bindings during prophage induction. Results show that different mutants with different binding deficiencies of Cro_2_ from O_*R*_1 and CI_2_ from O_*R*_3 do not influence the transition point of lysogeny transferred to lysis(Fig 7).

**Figure 7:**
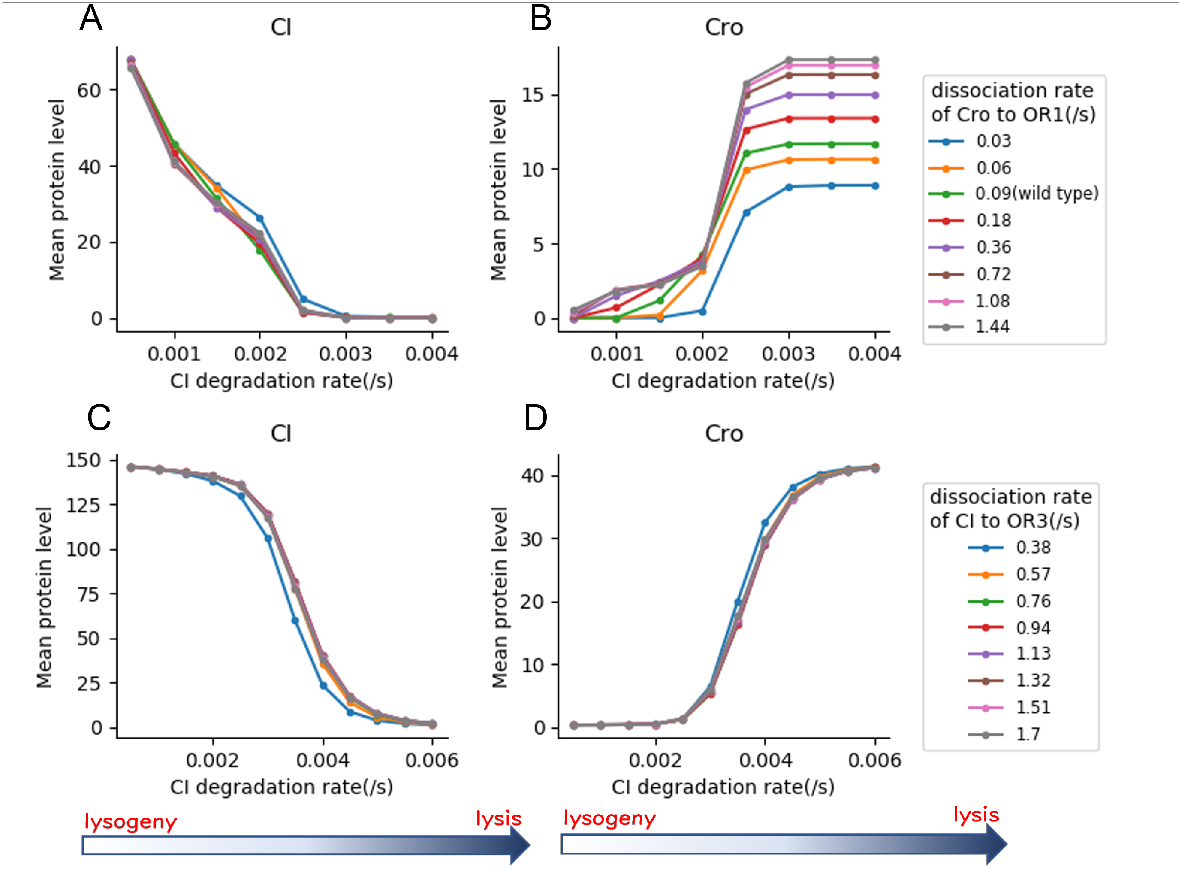
Cro_2_-O_*R*_1 and CI_2_-O_*R*_3 has little effect on the switch. K_*d*_ is CI_2_ degradation rate. (A-B) The mean copy number of CI protein(A) and Cro protein(B) under different CI_2_ degradation rate.different color represent different dissociation rate of Cro_2_-O_*R*_1.(C-D) The mean copy number of CI protein(C) and Cro protein(D) under different CI_2_ degradation rate. Different color represent different dissociation rate of CI_2_-O_*R*_3.

## 3 Discussion

In this study, we use a biochemical reaction network to characterize the gene regulatory module which control lysogeny-lysis switch of phage lambda. Using the method Accurate Chemical Master Equation (ACME), the steady state probability landscape of the system is directly computed. To explore the function of Protein Cro and CI in the system, we tested the influence of protein-DNA interactions by tuning their reaction rates. Furthermore, we explored under different UV irradiation rate, how protein-DNA interaction function in the lysogeny-lysis switch.

Our results show that Cro binding to OR3 facilitates switching from lysogeny to lysis and thus impairs lysogeny stability. Switch efficiency may be crucial for virus’ survival in a sense it could help virus lyse the host cell before the host die. This is consistent with previous experimental studies of Dodd’s and Svenningsen’s. Dodd’s experiments show when Cro only weakly represses PRM (through less binding to OR3), lysogens was induced in a smaller amount. In other word, the tighter Cro could binding OR3, more lysogens could be induced, which agrees with our results. The latter study of Svenningsen’s suggests that Cro protein may not to be important in lysogeny-lysis switch. However, from their results (Fig 3A), we observe the similar result as ours that lysogeny to lysis switch is delayed when Cro is deleted in the group OL cro-comparing with group OL cro+, which support our conclusion.

Our results also show CI binding to OR1 slows down the switching, therefore help maintain lysogeny state. Besides, our results suggest Cro binding to OR1 and CI binding to OR3 barely influence lysogeny stability during the prophage induction.

Overall, our work shows that Cro is important in lysogeny-lysis switch and explores the role of other protein-DNA complexes in the switch. By using method Accurate Chemical Master Equation (ACME), we could compute the exact steady state probability landscape of the system and further analyze dynamical behavior of it which allow us to dissect the function of proteins in the switch by tuning parameter such as reaction rate and CI degradation rate. The method ACME used here is very powerful and can be also used in other reaction networks.

## 4 Models and Methods

Our model describes the gene regulatory module as a reaction network. The species in the network include protein CI and Cro, operator right OR1, OR2, OR3 and protein-DNA complex where CI or Cro binding to these three operator sites. We assume the system is well-mixed with constant volume and temperature. The system is constructed by n molecular species x_1_, x_2_, …, x_*n*_ and m reactions with reaction constants r_1_, r_2_, …, r_*m*_. The k-th reaction is denoted as

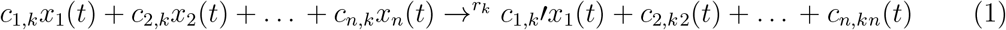

Where the microstate of the system at time t *x*(*t*) = (*x*_1_(*t*), *x*_2_(*t*), …, *x*_*n*_(*t*) ∈ ℤ^*n*^. The rate of reaction *R*_*k*_ that brings the microstate from *x*_*i*_ to *x*_*j*_ is denoted as

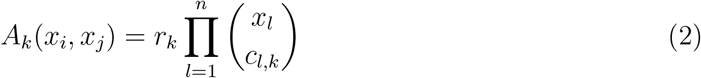

Where *r*_*k*_ is the reaction constant and *c*_*l,k*_ is the stoichiometry constant of the relevant reactants *x*_*l*_ in reaction *R*_*k*_

The discrete chemical master equation can be written using above definitions describing probability change of every microstates over time.

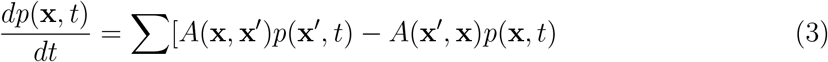

Where *p*(x, *t*) is probability of microstate x at time t. *A*(x, x^*′*^) is the transition rate from state x^*′*^ to state x. Using above equations, probability *p*(x, *t*) can be calculated using previously developed method Accurate Chemical Master Equation(ACME). Detail of the method can be found in ref [17].

## Notes

### Competing Interest Statement

The authors have declared no competing interest.

### Summary of Updates

author information updated.previous version has incorrect affliations

